# Pan-cancer association of DNA repair deficiencies with whole-genome mutational patterns

**DOI:** 10.1101/2022.01.31.478445

**Authors:** Simon G. Sørensen, Amruta Shrikhande, Gustav A. Poulsgaard, Mikkel H. Christensen, Johanna Bertl, Eva R. Hoffmann, Jakob S. Pedersen

## Abstract

DNA repair deficiencies in cancers may result in characteristic mutational patterns, as exemplified by deficiency of *BRCA1/2* and efficacy prediction for PARP-inhibitors. We trained and evaluated predictive models for loss-of-function (LOF) of 145 individual DDR genes based on genome-wide mutational patterns, including structural variants, indels, and base-substitution signatures. We identified 24 genes whose deficiency could be predicted with good accuracy, including expected mutational patterns for *BRCA1/2*, *MSH3/6*, *TP53*, and *CDK12* LOF variants. *CDK12* is associated with tandem-duplications, and we here demonstrate that this association can accurately predict gene deficiency in prostate cancers (area under the ROC curve=0.97). Our novel associations include mono- or biallelic LOF variants of *ATRX, IDH1, HERC2, CDKN2A, PTEN,* and *SMARCA4,* and our systematic approach yielded a catalogue of predictive models, which may provide targets for further research and development of treatment, and potentially help guide therapy.

## INTRODUCTION

The DNA damage response (DDR) and repair pathways are central to the genetic integrity of cells, and deficiencies may cause mutational patterns genome-wide^1–3^. Some DNA repair deficiencies are known to modulate the response to therapies: *BRCA1/2* deficiency renders cancers susceptible to treatment with PARP inhibitors^4^, mismatch repair-deficient cancers are sensitive to checkpoint inhibitors^5^ but resistant to alkylating agents such as temozolamide^6^, and *CDK12*-mutated cancers have a suggested sensitivity to CHK1 inhibitors^7^. Because of this, efforts have been made to annotate inactivating mutations in DDR genes^8^. However, the approach is limited by the lack of functional impact annotation of most variants, which are generally denoted as “variants of unknown significance” (VUS). Moreover, loss of gene activity could also occur by other means, such as transcriptional silencing.

A complementary approach is to investigate whether DNA repair deficiencies can be identified by DNA mutational patterns, also referred to as ‘mutational scars’. This approach has been pioneered for homologous recombination deficiency caused by *BRCA1/2* deficiencies (-d), which can be successfully predicted by measuring the accumulation of small deletions with neighbouring microhomologous sequences^2, 9, 10^, such as done by the HRDetect algorithm by Davies *et al.*^9^. The association with microhomologous deletions is due to the use of microhomology-mediated endjoining (MMEJ) to repair double-strand breaks in homologous recombination (HR) deficient tumours^11, 12^. Likewise, mismatch repair-deficiency causes an elevated rate of mono- and dinucleotide repeat indels genome-wide, a genetic phenotype denoted microsatellite instability^13, 14^. Mutations in other DNA repair genes have also been associated with mutational patterns, including the tumour suppressor gene *TP53*, which is associated with increased structural rearrangements and whole genome duplications^15, 16^ *CDK12* which is associated with a genome-wide phenotype of large tandem duplications^17, 18^.

The scope of this approach can now be evaluated systematically across DDR genes by exploiting available whole cancer genomes from thousands of patients^19, 20^.

To achieve this, mutations observed genome-wide may be condensed into mutational summary statistics for predictive modelling, including statistics based on single-base substitutions (SBS), indels, and different types of structural variants (SV). The SBSs are statistically assigned to so-called SBS signatures that are catalogued and enumerated within the COSMIC database^21^.

Some of these are associated with specific DNA repair deficiencies as well as genotoxic exposures, such as ultraviolet (UV) light and smoking. Each SBS signature captures the relative frequency of the different mutation types and their flanking nucleotides^22^.

Here we performed a systematic screen for DDR gene deficiencies that can be predicted through their association with genome-wide mutational patterns. We developed a generic approach to train predictive statistical models that identify associations with individual mutational summary statistics that capture the mutational patterns, including SBS signatures, indels, and large SVs. We applied it to 736 DDR gene deficiencies, considering both monoallelic and biallelic loss of function (LOF), identified across 32 cancer types, in a combined set of whole cancer genomes from 6,065 patients^19, 20^. The underlying aim was to identify novel associations with potential biological relevance and to evaluate whether DDR deficiencies can be predicted with sufficiently high certainty to have a potential for clinical application.

Our analysis revealed 24 DDR genes where deficiencies are associated with specific mutational summary statistics in individual cancer types across 48 predictive models. These results recapitulated the expected associations between mutational patterns and deficiencies of *BRCA1/2, TP53, MSH3/6,* and *CDK12*. We supplemented this knowledge by providing a predictive model of *CDK12* deficiency that achieved high accuracy (area under the receiver operator characteristic (AUROC) = 0.97) in prostate cancer. Furthermore, we present unexpected predictive models of several DDR deficiencies; *ATRX* and *IDH1* deficiency in cancers of the central nervous systems, *HERC2* and *CDKN2A* deficiency in skin, *PTEN* deficiency in cancers of the central nervous system and uterus, and *SMARCA4* deficiency in cancers of unknown primary.

## RESULTS

### DDR gene deficiencies across 6,065 whole cancer genomes

We compiled and analysed 2,568 whole genome sequences (WGS) from The Pan-Cancer Analysis of Whole Genomes (PCAWG)^19^ and 3,497 WGS from the Hartwig Medical Foundation (HMF)^20^. In total we investigated 6,065 whole cancer genomes of 32 cancer types (**Fig. 1a; Additional file 1, Supplementary Table 1**).

**Figure 1:**
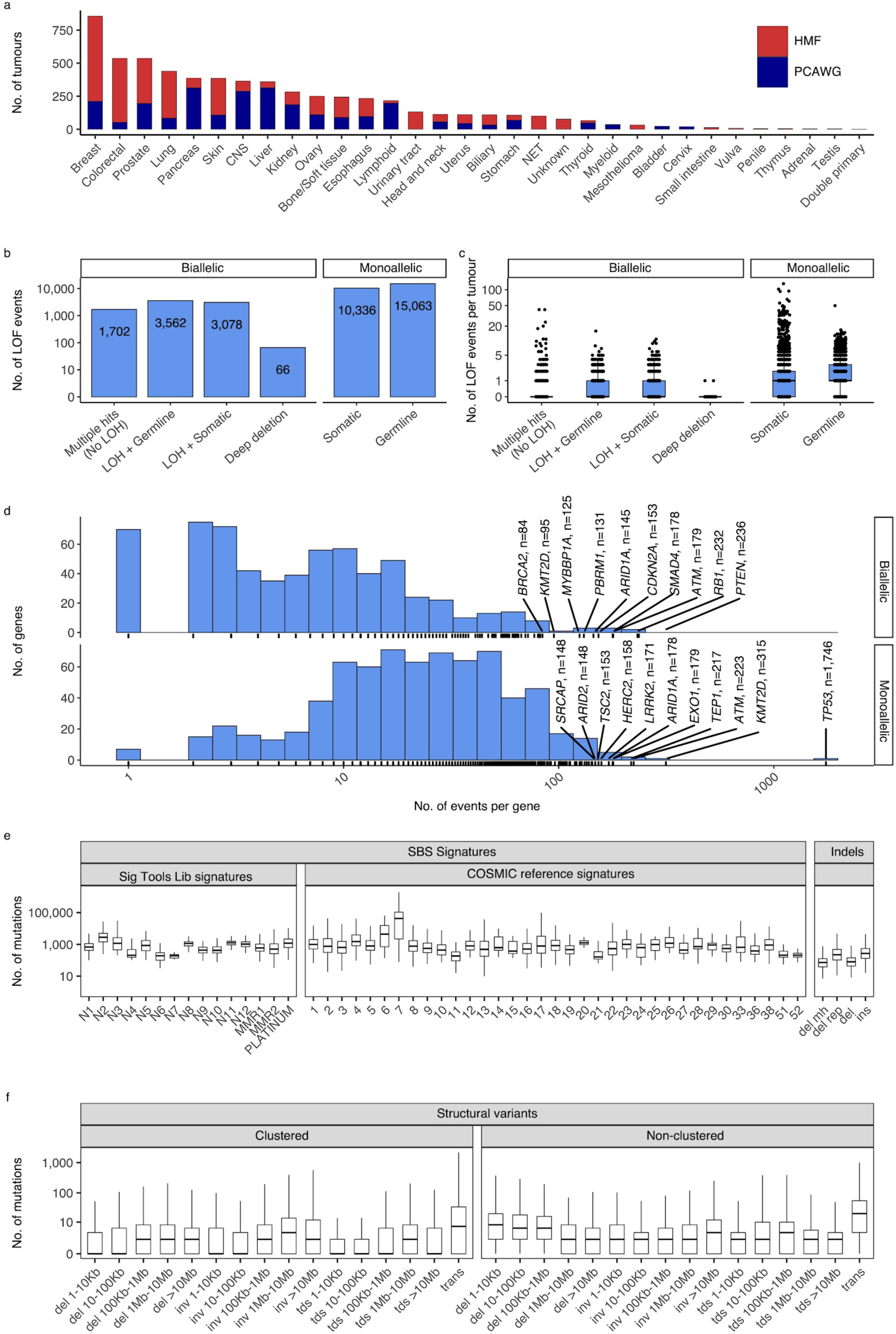
Cancer-types, DDR gene deficiencies, and mutational patterns. **a** Cohort sizes for the 32 cancer types comprising the 6,065 whole cancer genomes collected from the Hartwig Medical Foundation (HMF; n=3,497) and the PanCancer Analysis of Whole Genomes (PCAWG; n=2,568). **b** mono- and biallelic loss-of-function (LOF) events were annotated across 736 DDR genes based on both pathogenic variants and copy-number-losses (loss of heterozygosity; LOH), overall and (**c**) per patient (**d**) with varying no. of LOF events per DDR gene (x-axis; logarithmic). **e** Whole genome mutational patterns were represented as summary statistics and used as input features for the predictive models of DDR gene deficiency. Concretely, each patient was annotated with the number of single-base-substitutions (SBS) that are accounted to each SBS signature (Alexandrov *et al.*^22^; Degaspari *et al*.^27^), number of indels divided by context (mh = microhomology; rep = repetitive), and (**f**) number of structural variants divided by clusterness, size, and type (del = deletion; inv = inversion; tds = tandem duplication; trans = translocation)

For each genome, we evaluated 736 known DNA damage response (DDR) genes for both germline and somatic loss of function (LOF) events^23–25^. We annotated both mono- and biallelic loss-of-function (LOF) events, where each event could be either a single nucleotide variant, an indel, or a loss-of-heterozygosity (LOH) (**Fig. 1b,c; Additional file 1, Supplementary Table 2**). Pathogenicity of single base substitutions (SBSs) and indels was evaluated using a combination of CADD scores (>25; 0.3% most pathogenic variants) ^26^ and ClinVar annotation, when available (**Methods**).

We inferred a total of 8,408 biallelic DDR gene deficiencies, primarily through a combination of somatic or germline variants (SBSs and indels) with pathogenic potential (n=1,702), or LOH events combined with a single pathogenic germline (n=3,562) or somatic (n=3,078) variant (SBS or indel; **Fig. 1b**). On average we observed a single, biallelic DDR gene loss per patient, with some tumours showing extreme rates of somatic pathogenic mutations (**Fig. 1c**). As expected, *TP53* deficiency (*TP53*-d) was the most frequent LOF event (81 biallelic and 1,746 monoallelic events; 29% of tumours affected; **Additional file 1, Supplementary Table 2; Fig. 1d**), while 70% of DDR genes had biallelic deficiency in less than ten tumours across all cancer types (511/736; **Fig. 1d**). Among monoallelic events, we identified 15,063 pathogenic germline (59%) and 10,336 somatic (41%) events.

### Whole genome mutational patterns

We collected mutational summary statistics for each cancer genome, which were used as features for the downstream predictive models (**Fig. 1e,f; Additional file 2, Supplementary Table 3-4**). For SBSs, we evaluated exposure towards predefined sets of cohort-specific SBS signatures^22, 27^. Short indels and SVs were simply categorised and counted: Deletions were sub- categorised based on surrounding sequence repetitiveness and presence of microhomology.

SVs were sub-categorised by type (tandem duplications, inversions, deletions, and translocations), five size ranges (not relevant for translocations), and cluster presence (**Methods**). Several SBS signatures as well as some types of indels have suggested etiologies (collected in **Additional file 1, Supplementary Table 5)**.

### Statistical modelling of DDR gene deficiencies

For the downstream statistical analysis, we restricted our focus to DDR genes in cancer types with more than five biallelic LOF events in either PCAWG or HMF (n=194) or more than ten monoallelic LOF events (n=341) (**Additional file 1, Supplementary Table 6**). Using *BRCA2-*d in the set of HMF breast cancer tumours (n=645) as an example, we observed biallelic LOF events in 17 (2.6%; 14 germline, 3 somatic) tumours and monoallelic LOF events in 7 (1.1%; 4 germline, 3 somatic) tumours (**Additional file 1, Supplementary Table 8**). We further observed variants of unknown significance (VUS) events in 53 tumours (8.2%; 42 germline, 11 somatic), which were excluded from the analysis. The remaining *BRCA2* wild-type (WT) tumours (n=568; 88.1%) were used as a background set for training the predictive models (**Fig. 2a)**. The high fraction of germline pathogenic variants diminishes the probability of a reverse-causal relationship between the loss of *BRCA2* and the associated mutation patterns.

**Figure 2:**
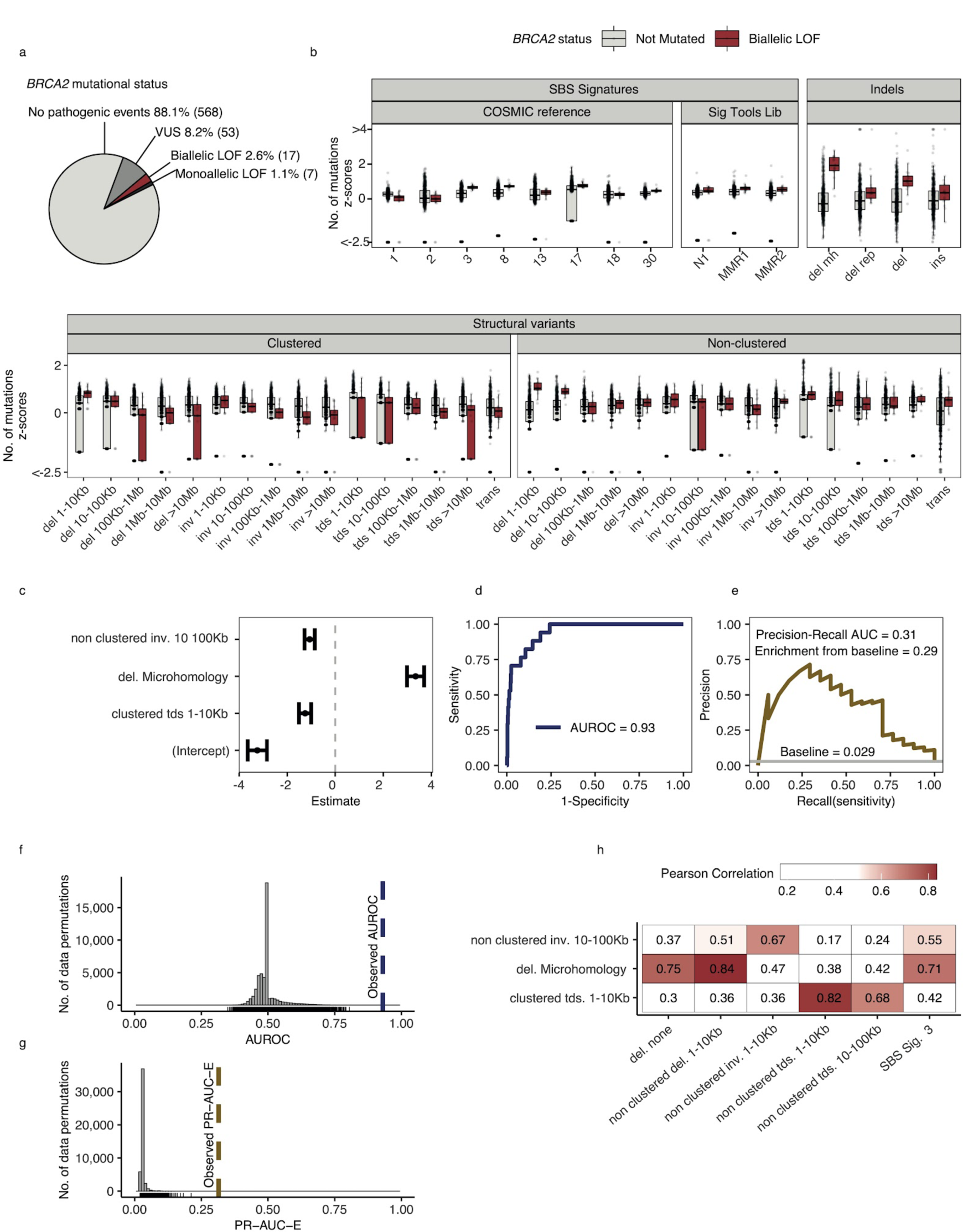
Predictive modelling of BRCA2 deficiencies in the HMF breast cancers. **a** Mutational status of *BRCA2* across 645 HMF breast cancer patients. **b** Mutational summary statistics for the HMF breast cancer patients divided by biallelic *BRCA2* loss-of-function (LOF; red) and *BRCA2* wild type (WT; grey) (selected predictive features in bold). **c** Predictive features and their coefficients for model of biallelic *BRCA2* loss with predictive performance measured in (**d**) AUROC and (**e**) PR-AUC (PR-AUC-E=PR-AUC - baseline=0.29; **Methods**). **f** Distributions of AUROC and (**g**) PR-AUC-E values obtained from 30,000 random data permutations compared to observed values (punctuated lines). **h** Correlation between selected predictive features (horizontal) and other highly correlated (Pearson corr. >0.65) mutational features (vertical).

For each of the 535 groups of tumours we trained a LASSO regression model and evaluated the ability to discriminate between deficient and WT tumours (**Methods**). For *BRCA2*-d we observed a strong association with the number of deletions at sites of microhomology (**Fig. 2b**), with a median of 608 deletions per patient in *BRCA2*-d breast cancers versus 81 in *BRCA2* WT breast cancers, in agreement with prior findings^2, 9, 10^. The LASSO regression also included non-clustered inversions 10-100Kb and clustered tandem duplications 1−10Kb, although both show high variance among tumours for both deficient and WT (**Fig. 2b**) and have considerably smaller coefficients, ultimately contributing little influence on overall predictive performance (**Fig. 2c**).

Notably, some models include features with negative coefficients. The biologically interpretation would be that tumours with a certain gene deficiency have fewer mutations attributed to a particular mutation pattern. Negative features were excluded in the development of the HRDetect algorithm^9^, but we include them as we cannot rule out the possibility that a DDR deficiency protects from specific types of mutagenesis. Though not distinguishable in this study, we suggest that negative coefficient features may derive in three ways: First, they may stem from enhanced repair; second, they may stem from the decomposition of mutation counts into signatures; and third, the mutated tumours may represent a subclass of patients in terms of age, gender, or tumour subtype with specific mutational patterns.

### Evaluating model performance

For each model, we evaluated the predictive performance using the area-under-the-receiver-operating-curve (AUROC) score as well as the precision-recall area-under-the-curve (PR-AUC) score. The PR-AUC score is a more robust measure for unbalanced data sets ^28^; however, the expected value for non-informative (unskilled) models equals the fraction of true positives and thus varies between models. Therefore, we used the PR-AUC enrichment over the true-positive rate (PR-AUC-E) as our selection criteria for predictive models. For the downstream analysis, we included (shortlisted) models with PR-AUC-E that was substantial (>0.2; more than two standard deviations above the mean across all 535 models) and significant (Benjamin-Hochberg false discovery rate, FDR<0.05; Monte Carlo simulations) (**Methods**; **Additional file 1, Supplementary Table 7**). In addition, we tested each model in the same cohort in the other data set whenever one or more tumours had mono- or biallelic loss of the gene. Due to the difference in biology between the two sets, we did not use this to explicitly evaluate the predictive performance of models but have included the PR-AUC-E values from the tests in **Supplementary Table 6 and 7**.

In the example of *BRCA2*-d in breast cancers of the HMF data set, our model achieved an AUROC of 0.93 and a PR-AUC-E of 0.29 (**Fig. 2d,e; Additional file 1, Supplementary Table 7**). Out of 30,000 permuted LOF-sets, none had a similar or higher PR-AUC score and we considered the model significant with a p-value < 3x10^-5^ (FDR adjusted q-value < 6x10^-4^) (**Fig. 2f,g**). The model achieved a PR-AUC-E of 0.37 when tested on the PCAWG data set, suggesting that the model may generalise across both metastatic and non-metastatic tumours.

This was further supported by the independent discovery of a similar model in the PCAWG data set, which had an almost similar predictive power in the HMF data (PR-AUC-E = 0.19; **Supplementary Table 7**). Notably, the *BRCA2*-d model did not include non-clustered deletions <100Kb, SBS signature 3, and SBS signature 8, all features which have been associated with BRCAness^9^. However, SBS signature 3 and non-clustered deletions 1-10Kb are included in the model when the deletions at sites of microhomology are omitted from the input data set, suggesting that they are excluded during feature selection due to high positive correlation with the number of deletions at sites of microhomology among HMF breast cancers (Pearson corr. > 0.7; **Fig. 2h**; **Additional file 1, Supplementary Table 9**).

Our selection criteria resulted in 48 shortlisted predictive models across 24 DDR genes (**Fig. 3a; Additional file 1, Supplementary Table 7)**. As exemplified for *BRCA2*, each model is specified by a set of predictive features representing mutational patterns associated with DDR gene LOF. We divided the models into four groups based on aetiology and origin: models of *BRCA1/2*-d (eight models of *BRCA2*-d and a single model of *BRCA1*-d; **Fig. 3b**); models of monoallelic *TP53*-d (11 models; **Fig. 3c**); models of various monoallelic gene deficiencies derived from colorectal cancer patients (eight models; **Fig. 3d**); and models including other DDR genes and cancer types, including previously undescribed associations (20 models; **Fig. 3e**).

**Figure 3:**
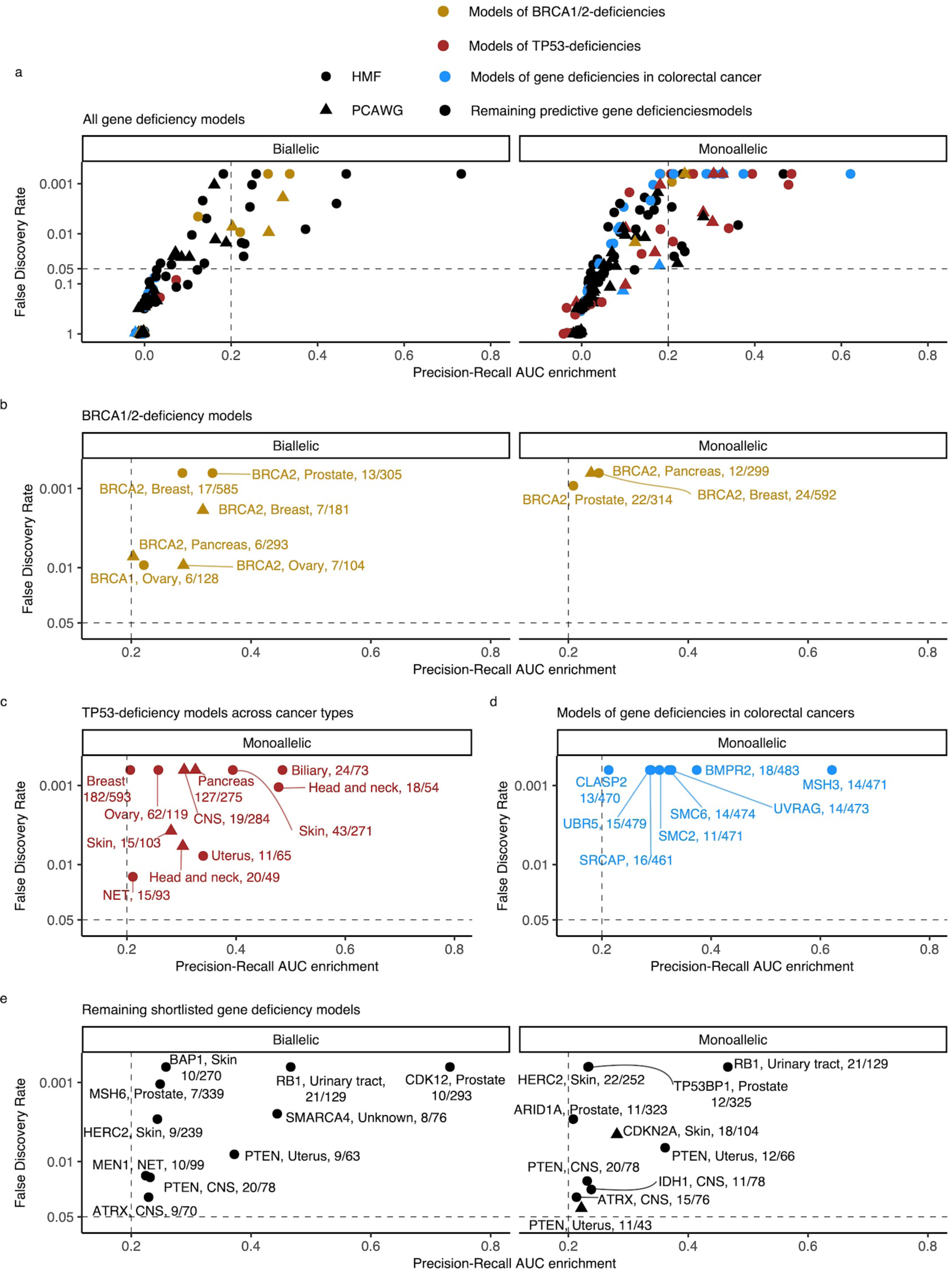
Predictive models of DDR gene deficiencies. **a** The precision-recall AUC enrichment PR-AUC-E; x-axis) and significance (FDR; logarithmic y-axis) of the 535 predictive models (one model per gene with more than five biallelic or more than ten tumours either mono- or biallelic mutated in either HMF or PCAWG in any one cancer type; **Methods**). Significance (q-value representing FDR) evaluated by counting equally or more-extreme PR-AUC-E values across >10,000 permuted data sets and applying Benjamini-Hochberg FDR control. Models with FDR below 0.05 and PR-AUC-E above 0.2 are shortlisted (**Methods**). **b** Shortlisted predictive models of deficiency of *BRCA1* or *BRCA2*; (**c**) *TP53* monoallelic predictive models; (**d**) monoallelic gene deficiency models across colorectal cancer patients; (**e**) and remaining gene deficiency models not contained in the other sub-groups. Numbers indicate the number of mutated out of the total number of tumours included in the development of each model.

### Recapitulation and predictive modelling of expected associations with *BRCA1/2* **deficiency**

Five models predicted biallelic loss of *BRCA2* in cancers of the ovary, prostate, pancreas, and breast. In addition, three models predicted *BRCA2* monoallelic loss in cancers of the pancreas, breast, and prostate. Finally, we derived a single model of biallelic *BRCA1* loss in ovarian cancer (**Fig. 3c; Fig. 4a**). All models significantly outperformed their Monte-Carlo simulations (q<0.05; Benjamini Hochberg FDR control) and had PR-AUC-E above 0.2 (**Fig. 4b,c**).

**Figure 4:**
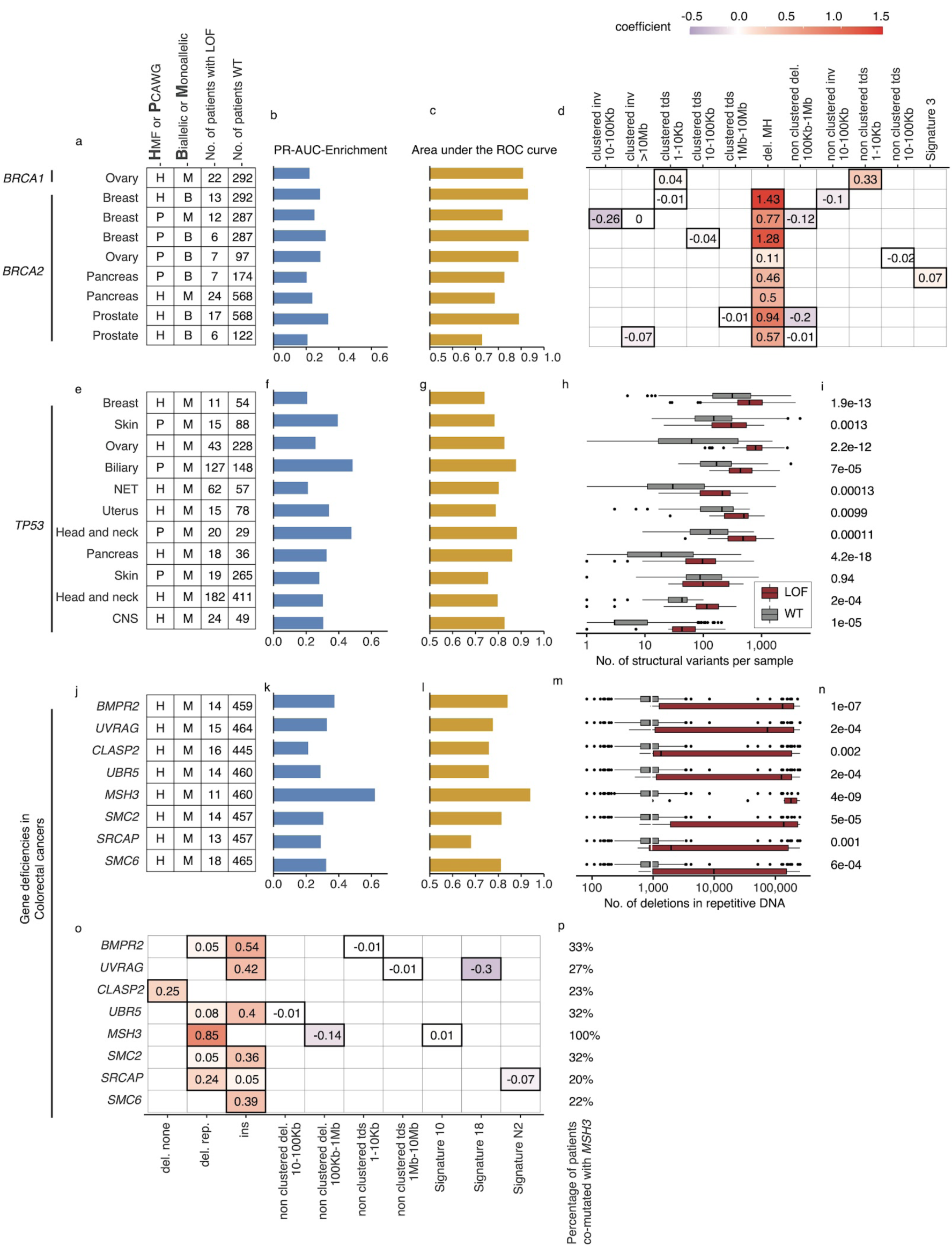
Predictive models with anticipated aetiology or origin. **a** Overview of predictive models for *BRCA1*-d and *BRCA2*-d, showing data source, type of model, and LOF-set statistics (**b**) PR-AUC-E, (**c**) AUROC, (**d**) and the predictive features and their coefficient for individual models. **e,f,g** Overview of predictive models of *TP53*-d (as in a-c). **h** For each cohort the number of structural variants (x-axis; logarithmic) for *TP53* LOF tumours (red) versus *TP53* wild type tumours (grey) and (**i**) the significance of their difference (two-sided Wilcoxon rank-sum test). **j,k,l** Predictive models of gene deficiencies in colorectal cancers (as in a-c). **m** Number of deletions in repetitive DNA (as in h) and (**n**) its significance (as in i)**. o** The predictive features of each model (as in d) and (**p**) the percentage of tumours that are co- mutated with *MSH3*.

All *BRCA2*-d models were predominantly predicted by deletions at sites of microhomology, consistent with the role of *BRCA2* in homologous recombination and suppression of microhomology-mediated end joining^29^. Both clustered and non-clustered tandem duplications in the range of 1 to 100Kb were included as features for various models, though with much smaller predictive power. This agrees with what was identified for *BRCA2* deficient tumours in prior studies^9, 10^. The biallelic breast cancer model based on PCAWG further included SBS signature 3 as a predictive feature (**Fig. 4d**). Contrasting to the models of *BRCA2*-d, *BRCA1-*d in ovarian cancer was exclusively associated with clustered and non-clustered tandem duplications (1- 10Kb; **Fig. 4d**). This aligns with prior studies^9, 10^, which also found *BRCA1*-d to be closely associated with a tandem-duplicator phenotype. In general, *BRCA1* and *BRCA2* were subject to predominantly germline pathogenic events, and not a single deletion at a site of microhomology, suggesting the expected forward causality (**Supplementary Table 8**). As for the loss of *BRCA2*, the model for loss of *BRCA1* loss in ovary had sufficient predictive power (PR-AUC-E = 0.3) in the other data set, suggesting that the model works independently of the metastatic capacity of the tumour (**Supplementary Table 7**).

### *TP53* deficiencies associate with increased numbers of structural variants

We detected 11 predictive models (four based on PCAWG and seven on HMF) of monoallelic *TP53*-d across cancers of the breast, skin, ovary, uterus, neuro-endocrine tissues (NET), biliary gland, head and neck, pancreas, and the central nervous system (CNS) (**Fig. 4e**). These predictive models performed with PR-AUC-E values ranging from 0.21 to 0.48 in breast and biliary gland cancers, respectively. Similarly, AUROC values ranged from 0.48 to 0.88, again in breast and biliary gland cancers (**Fig. 4f,g**). In line with existing literature^30^, *TP53-d* is associated with a significantly increased number of structural variants across the genome (Wilcoxon rank-sum test; **Fig. 4h,i**), except in skin cancers. The models of *TP53* loss in head and neck, skin, breast, and the biliary gland performed well (PR-AUC-E above 0.2) in the other data set, suggesting that the predictive performance generalises independent of metastatic tumour state (**Supplementary Table 7**).

### Colorectal cancer models derived from hypermutated mismatch repair deficient tumours

In the HMF colorectal cancers, we discovered eight predictive gene deficiency models (*MSH3, SMC2, SMC6, BMPR2, CLASP2, SRCAP*, *UBR5,* and *UVRAG)* of monoallelic LOF with PR-AUC-E ranging from 0.21 (*CLASP2*) to 0.62 (*MSH3*) (AUROC ranging from 0.68 for *SRCAP* deficiency to 0.94 for *MSH3* deficiency; **Fig. 4j,k,l**).

We suggest that the high number of models of monoallelic deficiencies may arise from spurious LOF events in DDR genes in a subset of colorectal cancers that are hypermutated. Some colorectal cancers are signified by mismatch repair deficiencies, such as LOF of *MSH3* or *MSH6*, creating a high number of deletions in repetitive DNA^13, 14^. Indeed, we found that this pattern was most profound among the *MSH3* mutated cancers (**Fig. 4m,n**). Furthermore, we found co-mutation with *MSH3* across the tumours underlying each model, ranging from 20% (*SRCAP*) to 33% (*BMPR2*) of the mutated tumours (**Fig. 4p**). This suggests that the models (except for the model of *MSH3*-d) might be the consequence of the hypermutator phenotype. In other words, the causality may be reversed in these cases, and the mutational process driven by *MSH3*-d may have caused the majority of their LOF events. This notion is supported by investigating the features of the models. All eight models are characterised by a single, primary predictive feature: Insertions (*SMC2, SMC6, BMPR2, UBR5, and UVRAG*), deletions in repetitive DNA (*MSH3, SRCAP*), or deletions not flanked by repetitive or microhomologous DNA (*CLASP2*) (**Fig. 4o**). Each of these features has a high positive correlation (Pearson corr. > 0.93) with the number of deletions in repetitive DNA. This correlation suggests that all eight models relate to a genome-instability phenotype, which may be driven by the *MSH3* co-mutated tumours or, potentially, a concurrent deficiency of other genes within the mismatch repair system (**Additional file 1, Supplementary Table 9**). Notably, the deficiency models of *UBR5, BMPR2, CLASP2,* and *SMC6* all had PR-AUC-E above 0.2 in the other data set, suggesting that these genes are associated with the mismatch repair phenotype regardless of metastatic state (**Supplementary Table 7**).

### Biallelic LOF of *MSH6* associated with increased number of deletions in repetitive DNA in prostate cancer

*MSH6*, a gene implicated in mismatch repair and microsatellite stability^14^, was mutated in both alleles in 7 out of 342 HMF prostate cancer patients. We observed pathogenic indels in *MSH6* in all 7 tumours, but only one of these in mono- or dinucleotide repeat DNA. *MSH6* deficiency could be predicted with high accuracy (PR-AUC-E=0.25; AUROC=0.98) by an enrichment of deletions in repetitive DNA (**Fig. 5a,b; Additional file 1, Supplementary Table 7**). This is consistent with existing findings of mismatch repair deficiency and its presence in metastatic prostate cancers^31^.

**Figure 5:**
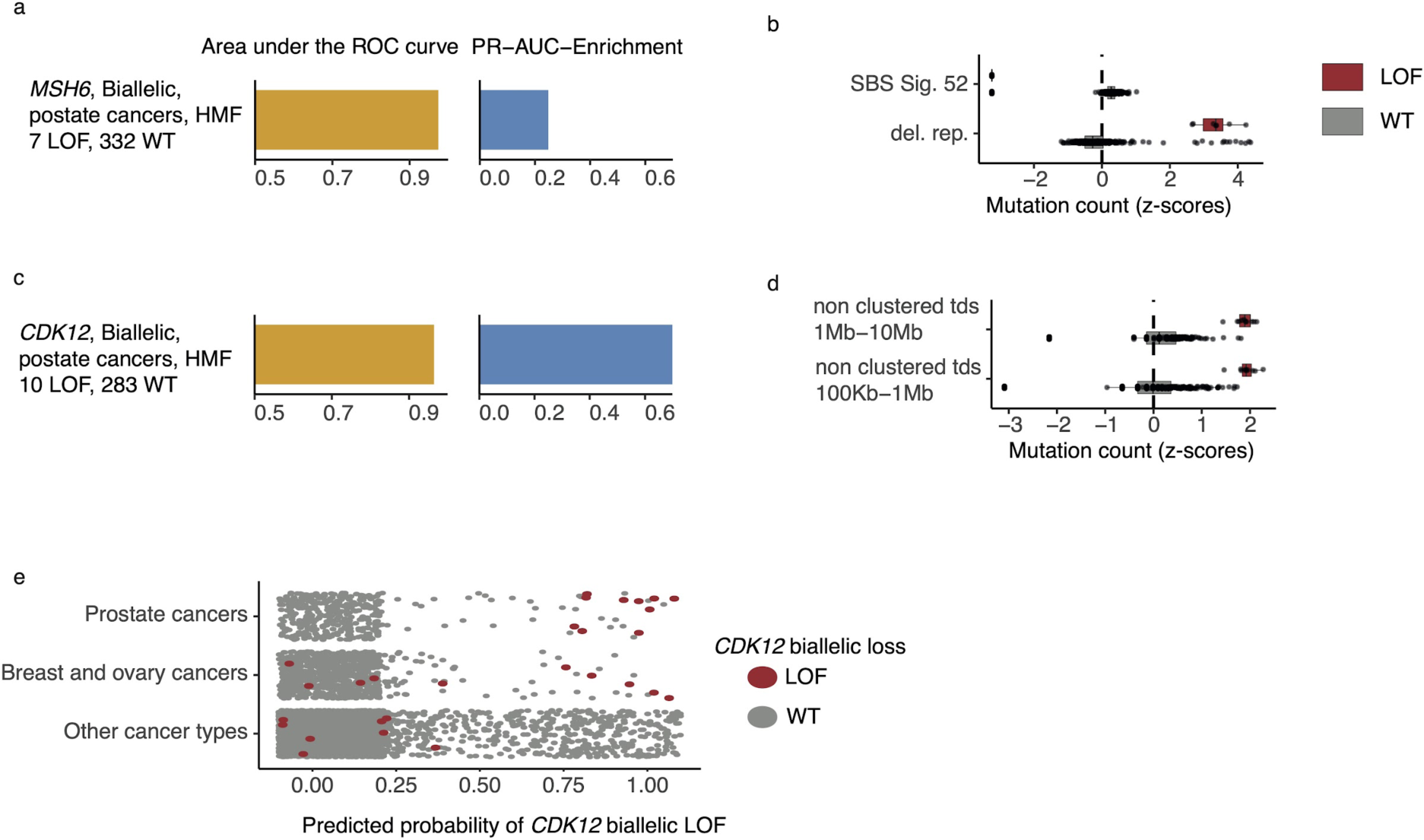
*CDK12* mutated prostate tumours are predicted by tandem duplications. **a** Biallelic predictive model *MSH6*-d in HMF prostate tumours and its PR-AUC-E and AUROC. **b** Boxplots of mutation counts between tumours that are *MSH6*-d loss-of-function (red) and *MSH6* wild-type (grey), (mutation counts are normalised and log-transformed; **Methods). c** Biallelic predictive model for *CDK12*-d with performance measures (as in a). **b** Boxplots of mutation counts for *CDK12* deficient and wildtype tumours (as in b). **e** *CDK12*-d predictive performance for different cancer types. Predicted probability of *CDK12*-d (x-axis) for tumours with *CDK12* LOF (red) and *CDK12* wild type (grey) are shown for prostate cancer, breast and ovary cancer, and all other cancer types (y-axis).

### High predictive power for *CDK12* deficiency

*CDK12* encodes a kinase that regulates transcriptional and post-transcriptional processes of the DDR^32–34^. We found that *CDK12*-d prostate cancers had an increased number of mid- and large-sized tandem duplications 100Kb - 10Mb, compared to CDK12-WT (**Fig. 5c,d**). Several studies have observed similar tandem duplication phenotypes in ovarian cancers ^17, 18, 35^ and castration resistant prostate cancers^36, 37^. In agreement, nine of the ten patients in our data set were treated with drugs associated with castration resistance (4 Enzalutamide, 3 Abiraterone, 1 Cabazitaxel,1 Pembrolizumab)^38^.

Whereas the tandem duplication phenotype has been previously associated with loss of *CDK12*, in this study we present the first predictive algorithm utilising and quantifying the high predictive value of these patterns (PR-AUC-E=0.73 and AUROC=0.97). Indeed, the loss of CDK12 has been demonstrated to sensitise cancer cells to CHK1-^7^ and PARP-inhibitors^39, 40^.

We went on to test the *CDK12*-d model across other cancer types (**Fig. 5c**). As expected, we observed predictive power in cancers of the ovary and breast, though at a lower level (PR-AUC- E=0.19 and AUROC=0.72). No predictive power was observed for the remaining cancer types. We only observed a single tumour with ballelic LOF of *CDK12* in PCAWG, but the predictive model was able to correctly identify this tumour, and reached a PR-AUC-E of 0.99 in the PCAWG data set.

### Novel predictive gene deficiency models

The shortlisted predictive LOF models further include biallelic LOF of *ATRX, PTEN, HERC2, MEN1, SMARCA4, BAP1,* and *RB1,* as well as monoallelic LOF of *ATRX, IDH1, PTEN, CDKN2A, ARID1A, TP53BP1, HERC2,* and *RB1* (**Fig. 3e; Additional file 1, Supplementary Table 7)**. The number of mutated tumours underlying each model varied from eight (biallelic *SMARCA*-d in cancers of unknown primary) to 22 (monoallelic *HERC2*-d in skin). We here present the models of *ATRX, IDH1, SMARCA4, CDKN2A, PTEN, and HERC2*; remaining models may be found in the supplementary material (**Additional file 1, 3, 4, Supplementary Table 7, Supplementary Fig. 1**).

### Predictive models of *ATRX*-d and *IDH1*-d in CNS cancers

We found that *ATRX-*d (monoallelic: PR-AUC-E=0.21, AUROC=0.71; biallelic: PR-AUC-E=0.23, AUROC=0.76; **Fig. 6a**) and *IDH1-*d (monoallelic: PR-AUC-E=0.24, AUROC=0.82; **Fig. 6b**) in CNS cancers were both predicted by a decreased number of SBS signature 8 mutations. In addition, *ATRX-*d was further predicted by non-clustered inv. 10-100Kb, although with a small coefficient and hence contributing limited discriminatory power. We discovered that 7 of11 *IDH1* mutated tumours were also *ATRX* mutated (28-fold enrichment compared to *IDH1* WT skin cancers; p=1.2x10^-8^, Fisher’s exact test) and that both *ATRX* and *IDH1* were predominantly hit by somatic mutations (14 of 15 for *ATRX*, 11 of 11 for *IDH1*).

**Figure 6:**
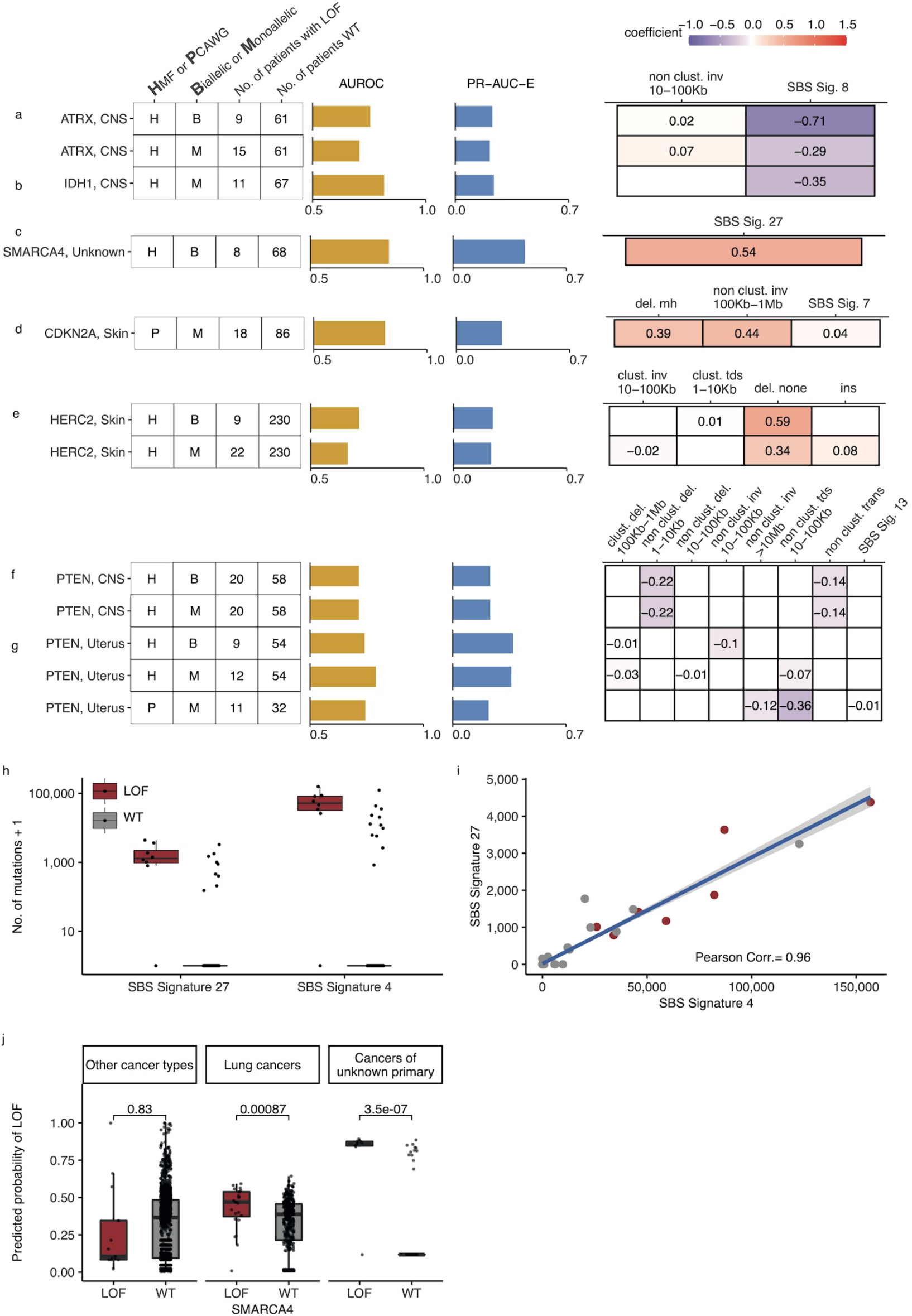
Novel predictive models of DDR gene deficiencies. **a** Predictive model of *ATRX*-d and its PR-AUC-E, AUROC, and selected features and their coefficients. Same information for predictive models of (**b)** *IDH1*-d, (**c**) *SMARCA4*-d, (**d**) *CDKN2A*-d, (**e**) *HERC2*-d, and (**f**) *PTEN-*d in CNS cancers and (**g**) uterine cancers. **h** Number of SBS sig. 27 and SBS sig. 4 (y-axis; logarithmic) among tumours of unknown primary with *SMARCA4* biallelic LOF (red) or wild-type (grey). **i** Pearson correlation between the per-tumour number (tumours of unknown primary; HMF) of SBS signature 27 (y-axis) and SBS signature 4 (x-axis; logarithmic) mutations, with an overlaid linear model (blue) and its 95% confidence interval (grey). **j** Using a model trained to predict *SMARCA4* biallelic LOF in HMF cancers of unknown primary, we evaluate the predictive power across cohorts (one-tailed Wilcoxon rank-sum test) displaying significant cohorts separately (colours as in **a**).

Co-mutation between *IDH1* and *ATRX* is well-described in gliomas^41^. Interestingly, LOF of either gene is associated with lack of SBS signature 8, a signature associated with BRCAness^9, 42^ and late-replication errors^43^. This suggests that CNS cancers with *IDH1/ATRX* deficiency are not subject to the same DNA lesions or repair processes as other CNS cancers and they may potentially belong to a separate patient subclass, though we could not identify evidence of this.

### Predictive model for *SMARCA4*-d in cancer of unknown primary

We discovered eight tumours (HMF) out of 77 with cancers of unknown primary with

*SMARCA4-*d (biallelic) that could be predicted with relatively high accuracy (PR-AUC-E=0.44; AUROC=0.85; **Fig. 6c**). These tumours showed an enrichment of SBS signature 27 (a signature first detected in myeloid cancers ^44^), which has been suggested to be a sequencing artefact though it also displays strong strand bias ^21^ (**Additional file 1, Supplementary Table 5**).

Among cancers of unknown primary, SBS signature 27 correlates strongly with SBS signature 4 (Pearson corr.=0.96), despite the signatures different composition (cosine similarity of signatures = 0.17) (**Additional file 1, Supplementary Table 9; Fig. 6h,i**). This suggests that SBS signature 4 may also be associated with *SMARCA4*-d and we indeed found that its predictive performance (PR-AUC-E of 0.43; AUROC=0.83) was almost equivalent to SBS signature 27. Signature 4 is associated with smoking across several cancer types^45, 46^; interestingly, *SMARCA4-*d is seen in aggressive thoracic sarcomas^47^ and strongly enriched among patients with a history of smoking^48^. Its gene product, BRG1, has been suggested as a lung cancer transcriptional regulator of genes that induce tumour proliferation^49^ and metastasis^50^. We evaluated the ability of SBS signature 4 to predict *SMARCA4*-d in other cancer types and found a significant predictive association in lung cancer, though much lower than for cancers of unknown primary (**Fig. 6j**). We also found a significant ability to predict *SMARCA4*-d by the number of SBS signature 27 mutations in cancers of the NET (Wilcoxon test, one-tailed p=0.002) and head and neck (p=4.8x10^-7^), but we could not evaluate this in lung cancers, as the signature is not among its set of cohort-specific signatures.

Twelve cancers had a high posterior probability of *SMARCA4*-d (**Fig. 6j**) despite being *SMARCA4* WT and lacking pathogenic events. No other single DDR gene was mutated among all twelve tumours, with *TP53* having the most LOF events (6 of 12 cancers). Given that cancers of unknown primary have disparate origins, this raises the possibility that the patients with high posterior probability of *SMARCA4*-d may have metastasized from a cancer type or subtype with both high levels of SBS signature 4 and 27, as well as high incidence of *SMARCA4*-d. Further studies are thus needed to clarify if the observed associations can be explained through such an ascertainment bias rather than causatively by *SMARCA4*-d.

### Predictive model for monoallelic *CDKN2A*-d in skin cancer

Both germline and somatic variants in *CDKN2A* are known to predispose for melanoma^51^. In the PCAWG skin cancer cohort, we found that a monoallelic predictive model of *CDKN2A*-d achieved relatively high accuracy (PR-AUC-E=0.28; AUROC=0.82). Its predictive features are enrichment of deletions at sites of microhomology, non-clustered inv. (100Kb-1Mb), and SBS signature 7, in order of predictive importance (**Fig. 6d**). Apart from the inversions, these features are also significantly associated with biallelic *CDKN2A*-d in the HMF skin cancer cohort (**Additional file 4, Supplementary Fig. 2a,b**) and are included in the corresponding predictive model, though it had lower predictive performance (PR-AUC-E=0.008; AUROC=0.542) and was not shortlisted (**Additional file 1, Supplementary Table 6**). In both HMF and PCAWG, most of the observed LOF events are somatic, with the majority being biallelic (16 of 18 in PCAWG; 32 of 33 in HMF; **Additional file 1, Supplementary Table 8**). The presence of deletions at sites of microhomology suggests a possible reduction in error-free double-stranded break repair in combination with an increased accumulation of SBS signature 7.

### Predictive model for *HERC2-d* in metastatic skin cancer

*HERC2*-d has been associated with susceptibility to developing melanoma^52^. We found *HERC2*- d predictable in HMF skin cancer patients (biallelic: PR-AUC-E=0.24, AUROC=0.73; monoallelic: PR-AUC-E=0.23, AUROC=0.66; **Fig. 6e)**, primarily based on enrichment of deletions in non-microhomologous and non-repetitive regions.

*HERC2* encodes a protein ligase that modulates the activity of P53^53^. We observed that seven of the nine tumours with biallelic *HERC2*-d also had a monoallelic pathogenic event in *TP53* (8- fold enrichment; p=4.6x10^-6^, Fisher’s exact test). Tumours that are co-mutated in *TP53* (monoallelic) and *HERC2* (mono- or biallelic) showed a significantly higher number of deletions compared to tumours with LOF in either gene alone (Wilcoxon test, One-tailed p<0.031) and cancers that are wild-type for both genes (p<7.2x10^-7^) (**Additional file 4, Supplementary Fig. 2c**).

### *PTEN* deficiency associates with fewer structural variants in CNS and uterine cancers

*PTEN* is a tumour suppressor gene found in various cancer types^54, 55^ and its deficiency was found to be associated with mutational patterns in uterine and CNS cancers (**Additional file 1, Supplementary Table 7**). In CNS, we acquired two identical models from the HMF data set, as we observed no monoallelic events without a second hit (mono- and biallelic PR-AUC-E=0.37, AUROC=0.74; **Fig. 6f**). In uterine cancer, we acquired significant predictive models from both HMF and PCAWG (HMF, biallelic: PR-AUC-E=0.37, AUROC=0.74; HMF, monoallelic: PR-AUC- E=0.36, AUROC=0.79; PCAWG, monoallelic: PR-AUC-E=0.22, AUROC=0.74; **Fig. 6g**). In addition, the models of *PTEN* loss in uterus had predictive power in the other data sets, suggesting signal robustness and independence of metastatic state (the PCAWG derived model had PR-AUC-E of 0.55 in the HMF data; the HMF derived model had a PR-AUC-E of 0.27 in PCAWG; **Supplementary Table 7**). The model of biallelic *PTEN*-d in uterine cancers is primarily based on depletion of non-clustered inv. 10-100Kb, whereas both the HMF and PCAWG models of monoallelic deficiency are primarily based on depletion of non-clustered tandem duplications (10-100Kb) (**Fig. 6g**). In contrast, our models of *PTEN*-d in CNS cancers from HMF were based on depletion of both non-clustered deletions 1-10Kb and non-clustered translocations (**Fig. 6f**).

## DISCUSSION

The cancer-specific, incomplete repair of endogenous and exogenous DNA lesions leave specific genome-wide mutational patterns. Their detection provides potentially powerful information on the fidelity of individual DNA repair pathways and response to chemo- and immunotherapies. Taking advantage of a large pan-cancer data set, our analysis shows that mutational patterns are associated with DNA repair defects across a wide range of cancers and repair mechanisms. In this study, we have contributed concrete initial predictive algorithms for mutational patterns of several DDR gene deficiencies, with potential use for clinical intervention.

The clinical scope of our approach is exemplified by recent regulatory approval for PARPi administration, which was supported by statistical predictions of homologous recombination deficiency (HRD) based on mutational patterns^56^ We have similarly predicted *CDK12* deficiency (*CDK12*-d) which has previously been associated with a tandem duplication phenotype– with power similar to that of HRD detection. This suggests a clinical benefit of clinical application of predictive algorithms to test for *CDK12*-d, specifically when considering the application of CHK1-inhibitors in prostate, ovary, and breast cancer treatment^7^.

We also find high predictive power for *MSH6*-d in prostate cancers by counting deletions in repetitive DNA, also known as the microsatellite instability phenotype (MSI)^57^. It is common practice to search for signs of MSI in colorectal cancers, endometrial cancers, and aggressive prostate cancers. This prediction is most commonly made using a panel of repetitive DNA regions, such as the Bethesda panel^58^. Here we demonstrated high predictive power in prostate cancers based on mutation summary statistics, which may be routinely extracted from whole genome sequencing. The correct identification of tumours with high MSI supports the administration of immune checkpoint blockade treatment, as the mutational phenotype leads to an increased expression of neo-peptides and thus an increased sensitivity towards the immune response^59, 60^.

Our systematic approach identified a similar model of *PTEN* LOF in uterine cancer in the PCAWG data and the HMF data; either model also having predictive power in the other data set. The re-discovery in either dataset grants further trust to this model, and further experimental investigation is warranted to understand the underlying aetiology of the mutation patterns, and the potential for clinical benefits.

Recent studies have suggested the use of CDK4/6 inhibition in treating *SMARCA4* deficient tumours^61^. We discovered that a large subset of cancers of unknown primary (>10%) have *SMARCA4* deficiency and that these cancers can be accurately predicted from their whole genome accumulation of SBS signature 4 or SBS signature 27 mutations. This is a clinically challenging cancer type due to the lack of a primary cancer to guide treatment and any such prediction may serve in selecting treatment for these complex cancers, given additional experimental evaluation.

Of note, the inclusion of monoallelic events allowed for larger sets of tumours with DRR gene LOF, thereby increasing the power of our study when causative associations with mutational patterns exist. This is for instance the case for *TP53,* which is known to be functionally impacted by monoallelic events^62^. We shortlisted 11 models of TP53-d, which were all monoallelic. We further found that inclusion of monoallelic LOF events matched the predictive power for *ATRX* biallelic LOF in CNS cancers and *HERC2* biallelic LOF in skin cancers, suggesting that single genetic hits are sufficient to affect repair and ensuing mutational patterns for these genes.

Incorrectly annotated LOF events in DDR genes may affect both the ability to discover associations with mutational patterns and the performance of any associated predictive models. On the other hand, highly stringent LOF criteria may result in true LOF events being overlooked, and create small deficiency sets with insufficient power to discover true associations. For the annotation of germline LOF events, we restricted our focus to rare variants (population frequency ≤0.5%), as more common variants are likely benign. Given the high number of variants evaluated overall, false positive germline LOF calls are expected. However, they are unlikely to associate with specific mutational patterns and thus are unlikely to contribute significant, shortlisted false positive predictive models of DDR deficiency. Likewise, in cancers with hypermutator phenotypes, somatic LOF events may be caused by and associated with specific mutational patterns. This leads to predictive models that in effect detect instances of reverse causality, as discussed for the monoallelic models found in colorectal cancer. We investigated whether reverse causality was a likely explanation for an association captured by a predictive model by evaluating whether the annotated LOF events matched the predictive mutational features (**Additional file 1, Supplementary Table 9**). In general, biallelic LOF criteria is less sensitive to wrongly annotated LOF events than monoallelic criteria, as double hits are rare compared to single hits. For both mono- and biallelic predictive models, the data permutations ensure that the observed association between the DDR gene LOF events and given mutational patterns are surprising.

Our conservative LOF curation, and a lack of expression or protein level data, is expected to cause some LOF events to be missed. This may explain a high posterior probability of a particular gene LOF in some tumours, but no evidence of genetic disruption (as seen in **Figure 5e** and **Figure 6h**). This highlights the scope of mutation-pattern-based predictions, in particular in tumours without canonical DDR gene LOF events. At a very least, such tumours could be considered for further scrutiny for LOF of the individual DDR gene that they are predicted to have lost.

Our systematic approach provides a proof-of-concept that will become increasingly powerful as the available data sets increase in number and size. For the current data sets, consistent validation of detected associations was challenging due to small cohorts and differences in cancer biology. To establish the basis for any future synthetic-lethality uses of our predictive models, it would be desirable to establish causal relationships. This has been beyond the scope of this study but could be achieved by whole-genome-sequencing of cell-lines or organoids with individual DDR gene knockout. Alternatively, the loss of DDR genes can be explored in animal models as previously done for several DDR genes including *BRCA1/2*^63^, MMR genes^64, 65^, and *TP53*^66^.

The current study yields a catalogue of predictive models that captures both known and novel associations between DDR gene deficiencies and mutational patterns. The included DDR genes may provide targets for research and development of treatment. With further optimization, predictive models such as these may guide the selection of therapy by adding certainty of disabled or compromised repair deficiency phenotypes found in cancers of individual patients.

## Materials and methods

### Data

The analysis was conducted on 6,098 whole cancer genomes and included a set of 6,065 whole cancer genomes after filtering (see below). The data came from two independent data sets: the Pan-Cancer analysis of Whole Genomes (PCAWG; tumours = 2,583; ICGC study ID. EGAS00001001692)^19^ and the Hartwig Medical Foundation (HMF; tumours = 3,515; Acc. Nr. DR-044)^20^. The data sets contain tumours from 32 cancer types and represent diverse patient groups in terms of age, gender, and disease history. PCAWG consists of tumours both with and without metastasis, whereas HMF exclusively consists of donors with tumours showing metastatic capability. To best relate the two data sets, all tumours are catalogued by the site of primary disease. For 77 of the metastatic HMF tumours, the primary site was unknown, and these are annotated as such (**Figure 1a; Additional file 1, Supplementary Table 1**).

### Curating samples

In the HMF data, we selected the earliest available sample whenever multiple samples existed for the same patient. For the PCAWG data, we used the official whitelist, obtained at the ICGC resources (https://dcc.icgc.org/pcawg), and for patients with more than one sample, we always selected the earliest, whitelisted sample available. We discarded six PCAWG samples of bone/soft tissue, as these were discarded in prior publications based on the PCAWG data set^27^ leaving 2,568 PCAWG samples and 3,497 HMF samples across 32 sites of the body. A list of all Donor IDs, Sample IDs, and primary sites of disease may be found in **Additional file 1, Supplementary Table 1**.

### Curating variants for loss-of-function annotation

We annotated variants across a set of 736 genes related to the DNA damage response (**Additional file 1, Supplementary Table 2**). The set of genes is combined from three sources, Knijnenburg *et al*.^24^; Pearl *et al*.^23^; and Olivieri *et al*.^25^.

We filtered the downloaded variant call files (VCF) of all samples for variants between the Ensembl (GRCH37/hg19) start- and end position of each DDR gene (see coordinates in **Additional file 1, Supplementary Table 2**). We included variants classified as PASS in the VCF files and variants of the PCAWG data set supported by at least two of our four variant callers. Furthermore, we discarded all variants which occurred in more than 200 samples across the two data sets in order to avoid noise arising from single nucleotide polymorphisms (SNPs) called as single nucleotide variants (SNVs), SNPs with high frequency in particular populations and possibly technical artefacts. We also discarded somatic variants with variant allele fractions (VAF) below 0.2 and variants where gnomAD (V2.1.1) showed a germline population frequency above 0.5%^67^.

### Annotating pathogenic variants and mutations

We annotated all variants and mutations with CADD phred scores (V1.6) in order to separate likely pathogenic variants from likely benign variants^26^. All variants with CADD phred scores of 25 or higher were considered pathogenic, whereas non-synonymous mutations with CADD phred scores below 25 and above 10 were considered variants of unknown significance (VUS). Variants with CADD phred scores below 10 were considered benign. A CADD phred score of 25 is a conservative threshold^68^, and only includes the 0.3% most-likely pathogenic variants. In addition, we annotated all variants with their status in the ClinVar database (when present).

Combining ClinVar and CADD phred scores ensured that we would be able to discover associations across all DDR genes, not only genes with ClinVar annotations, which are expected to be incomplete for any given gene.

### Annotating loss of heterozygosity and deep deletions

We also used the copy number profiles of each sample to discover genes with biallelic or monoallelic losses of parts of genes. Any overlap between a gene and a copy number loss was indicated as a loss-of-function event, under the assumption that losing any part of the protein coding DNA is detrimental to the complete protein product. We considered events in which the minor allele copy number was below 0.2 to be a loss of heterozygosity (LOH), whereas we considered events with a total tumour copy number below 0.3 (major and minor allele summarised) as deep deletions. This cutoff is adapted from the work of Nguyen *et al*.^10^.

### Annotating biallelic and monoallelic gene hits

We considered genes with a single pathogenic variant, either somatic or germline, to be monoallelic hit (n = 25,399). In cases where the monoallelic hit is accompanied by a LOH, we considered the event a biallelic loss (n = 6,640). Finally, genes that were completely depleted in a sample were considered biallelic lost (n = 66). We did not consider a single LOH event as a monoallelic loss due to the broad impact and high frequency of such events, with ∼25 times higher rates of LOH than deep deletions (**Figure 1b,c**).

### Mutational patterns of single base substitutions, indels and structural variants

We summarised the genome-wide set of somatic mutations by using Signature Tools Lib, developed by Degaspari *et al.*^27^ Firstly, we used Signature Tools Lib to count single nucleotide variants by base change and context, and assigned these counts to a set of organ-specific signatures, as recommended by Degaspari *et al*. We then have converted the organ-specific signature exposures to reference signature exposures using the conversion matrix found in the supplementary material of the paper by Degaspari *et al.*^27^ For twelve signatures (N1-N12) this was not possible, and these signatures are considered to be novel. Likewise, two signatures associated with mismatch repair (MMR) deficiency, MMR1 and MMR2, are associated with several of the COSMIC MMR signatures. For these, we preserved the signature label as given by Signature Tools Lib. For cohorts with no organ-specific signatures (cancers of unknown primary, neuroendocrine tissue, thymus, urinary tract, penis, testis, small intestine, vulva, double primary, and the adrenal gland), we have assigned mutations directly to the full set of 30 reference signatures (COSMIC signatures 1-30). Note that we excluded age-related signature 1 from the modelling, as this signature confounded by acting as a proxy for the age of the patient so that the model would be learning to differentiate old from young patients rather than patients with activity of different mutational processes or repair deficiencies.

Secondly, Signature Tools Lib was used to count indels: insertions were simply counted, whereas deletions were separated by the DNA context of the deletion, being microhomologous, repetitive, or none of the two. Microhomology was defined by whether the deleted sequence was similar to the region immediately 3’ of the breakpoint, as this indicates repair by microhomology-mediated end joining. Repetitive deletions were defined by whether there is a repeat of the indel at the 3’ end of the breakpoint^27^. We decided to not further compose indels into signatures, for the ease of interpretation; the type of deletion has a clear relation to the mechanism of repair, which is what we are investigating in this paper.

Finally, we used Signature Tools Lib to count structural variants. SVs were separated into two groups, clustered and non-clustered based on the average distance between rearrangements (the inter-rearrangement distance). Regions with at least 10 breakpoints having at least 10 times smaller inter-rearrangement distance than the average across the genome of that cancer, were considered clusters^69^. SVs were further divided based on the type of mutation: Deletions, tandem duplications, inversions, and translocations. Deletions, tandem duplications, and inversions were further divided by size, with intervals being 1-10Kb, 10-100Kb, 100Kb - 1Mb, 1- 10Mb, and finally mutations with lengths above 10Mb. As with indels, we decided not to decompose SV counts into rearrangement signatures, to ease interpretability and because SV signatures are less established in the field. The per-sample exposure to each summary statistic may be seen in **Additional file 1, Supplementary Table 3** (log transformed and scaled in **Additional file 1, Supplementary Table 4;** see method below) and the suggested aetiology (when existing) may be seen in **Additional file 1, Supplementary Table 5**^21, 42^.

### Preparing subsets of data for modelling

We separated the two data sets and divided the tumours into their respective cancer-type cohorts. We stratified the data for each DDR gene; designating whether the patient had a biallelic pathogenic variant, monoallelic pathogenic variant, or no pathogenic variants. We excluded tumours with inconclusive variants (CADD phred >10, <25) within the gene. For modelling of cancers with biallelic variants in a gene, we selected all DDR gene-cohort combinations where more than five tumours had biallelic variants in the same gene (n=194. Likewise, we selected all cases with more than ten tumours with monoallelic variants (n = 341) (**Additional file 1, Supplementary Table 6**). This setup means that each sample may be included in developing several models but occur only once in each model as either mutated or non-mutated. Biallelic mutated tumours were also included in the models of monoallelic loss, as we consider biallelic loss a special case of monoallelic loss.

### Generating models using LASSO regression

For each model we calculated per-sample weights to counter the imbalance in the data, so that the weight of each sample was one minus the proportion of tumours with this mutational status (either pathogenic or non-pahogenic/wildtype). We then used logistic regression model with least absolute shrinkage and selection operator (LASSO) regularisation (henceforth LASSO regression), with the alpha parameter at 1, from the R-package “glmnet” (v4.0)^70^ to select features with predictive power. The LASSO regression was selected to achieve a sparse set of associated features that could be readily interpreted. This is in line with the methodology used in HRDetect by Davies *et al.*^9^.

For each of k cross-fold validation sets (with k being the number of biallelic or monoallelic hit tumours, respectively), we ran a LASSO regression with a nested 5-fold cross-validation and the assigned weights (as mentioned above to counter imbalance in the data), to produce a set of lambda values. From the LASSO regression we selected the lambda value corresponding to the binomial deviance which is one standard deviation away from the observed minimal binomial deviance of the regression, in order to avoid overfitting (overfitting may occur if taking the lambda of the minimal binomial deviance)^70^. In a few cases, the LASSO regression converged to larger sets of features than what is justifiable by the low number of mutated tumours, and so we limited the number of features to one feature per ten mutated tumours, rounded up and added one (e.g. 12 mutated tumours gives basis for maximum three features). For each of the k cross-folds we used the model to predict the left-out data and used these predictions for the evaluation of the model performance.

Finally, we generated a model by including the entire data set. This was done to train the best possible model in terms of selected features and coefficients, and this is the model reported in the main figures, whereas the performance measures shown in the main figures are derived from the k-fold cross-validation.

### Evaluating model predictive performance

We used k-fold cross validation to get a measure of the predictive performance of each model. Due to the strong imbalance in the data, we use the precision-recall area-under-the-curve (PR-AUC) enrichment from the true-positive rate (PR-AUC-E) as our statistic of performance, while also reporting traditional AUROC scores. The true-positive rate is the baseline that would be expected for a non-informed model. By subtracting the true-positive rate from the PR-AUC of each model, we can compare the performance between models despite different sample sizes.

### Model selection

We selected models for further analysis (our shortlist) based on both their PR-AUC-E performance (effect size) and their significance. Across the 535 initial models, we observed a mean PR-AUC-E of 0.04 with a standard deviation of 0.099 (90%-quantile = 0.18) (**Additional file 1, Supplementary Table 6**). The PR-AUC-E threshold was set at 0.2 and thus roughly two standard deviations above the overall mean (**Additional file 1, Supplementary Table 7**).

To evaluate significance, we generated a null-distribution for each of the 535 initial models. This was done by permuting the mutation state of the underlying tumours and then running the LASSO regression on the permuted data. We ran a minimum of 10,000 permutations (n) for each of the 535 models, storing the PR-AUC-E for each permutation. For models where the number (r) of permutations leading to PR-AUC-E values as or more extreme than the original model was smaller than five, we ran an additional 10,000 permutations, up to a maximum of 30,000 permutations. A p-value (p=(*r* + 1)/*n*) was calculated for each model (**Additional file 1, Supplementary Table 6**). The Benjamini-Hochberg procedure was used to control the false discovery rate and resulting adjusted p-values (q-values) smaller than 0.05 were considered significant (**Additional file 1, Supplementary Table 7**).

In conclusion, 48 models of 24 genes were shortlisted for further investigation. Of these, we describe 43 in the main text and, more briefly, 5 in **Supplementary text** and **Supplementary figure 1.** The inclusion into the main manuscript was done by collegial evaluation of interest to the field and does not represent a definitive difference in quality of the models.

## DECLARATIONS

### Ethics approval and consent to participate

We analysed data generated and made available by the Pan-Cancer Analysis of Whole Genomes (PCAWG)^19^ Consortium of the International Cancer Genome Consortium (ICGC) and The Cancer Genome Atlas (TCGA) as well as the Hartwig Medical Foundation^20^. The research conforms to the principles of the Helsinki Declaration.

### Availability of Data and Materials

The data sets supporting the conclusions of this article are from the Pan-Cancer Analysis of Whole Genomes and from the Hartwig Medical Foundation. The public parts of the PCAWG data set is available at https://dcc.icgc.org/releases/PCAWG, whereas controlled files may be accessed through gbGaP and daco, as instructed on this site https://docs.icgc.org/pcawg/data/.

The ICGC study ID of the project is EGAS00001001692.

The HMF data used in this project may be found by accession code DR-044 and can be obtained upon request at the Hartwig Medical Foundation (https://www.hartwigmedicalfoundation.nl/en).

### Code availability

The code needed to reproduce the analysis will be made available at https://github.com/SimonGrund/DDR_Predict , including data pre-processing, modelling of the 535 predictive models and modelling of ≥10,000 data permutation null models for each of the 535 models.

## Author contributions

The project was conceived and designed by JSP and SGS, with early contributions by JB. The data analysis was carried out by SGS under supervision of JSP with input from MHC, including data preparation, exploratory analysis, and statistical modelling. SGS generated all figures and tables. SGS, AS, GAP, ERH, and JSP interpreted the results. SGS, ERH and JSP wrote the manuscript. All authors accepted the manuscript.

## Acknowledgements

We thank the Pan-Cancer Analysis of Whole Genomes (PCAWG), The International Cancer Genome Consortium (ICGC), and The Cancer Genome Atlas (TCGA) for access to whole cancer genomes. We would also like to thank the Hartwig Medical Foundation (HMF) and the Center for Personalized Cancer Treatment (CPCT) for creating and providing access to metastatic whole cancer genome data. We thank Jenny Gruhn and Amy V. Kaucher for valuable comments on the manuscript. We thank Dr. Andrea Degaspari for aid in using Signature Tools Lib for summarising mutation patterns.

## LIST OF ABBREVIATIONS

AUROC: Area under the receiver operator characteristic
CNS: Central nervous system
DDR: DNA damage response
HMF: Hartwig medical foundation
HRD: Homologous recombination deficiency
Indel: Insertion or deletion
LASSO: least absolute shrinkage and selection operator
LASSO regression: Logistic regression with LASSO regularisation
LOF: Loss of function
LOH: Loss of heterozygosity
MMR: Mismatch repair
MSI: Microsatellite instability
PCAWG: Pan-cancer analysis of whole genomes
PR-AUC: Precision-recall area under the curve
PR-AUC-E: Precision-recall area under the curve enrichment from baseline
SBS: Single base substitution
SV: Structural variant
UV: Ultraviolet
VUS: Variants of unknown significance
WGS: Whole genome sequence
WT: Wild type
-d: Deficiency

## Additional files

Additional file 1: Supplementary tables 1,2 and 4-9; Microsoft Excel (xlsx)

Additional file 1, Supplementary Table 1: All included tumours and their primary tumour locations

Additional file 1, Supplementary Table 2: 736 DDR genes, hg19 coordinates and the number of pathogenic events across 6,065 cancer genomes

Additional file 1, Supplementary Table 5: Proposed Etiologies of base substitution signatures Additional file 1, Supplementary Table 6: All models (n=535)

Additional file 1, Supplementary Table 7: Shortlisted models (n=48)

Additional file 1, Supplementary Table 8: Pathogenic events in each of the 535 LOF-sets Additional file 1, Supplementary Table 9: Correlation between features in shortlisted models

Additional file 2: Supplementary table 3-4; zip-compressed; tab-separated values (.tsv), may be opened in Microsoft Excel

Additional file 1, Supplementary Table 3: All SBS signature contributions, indels counts, and SV counts, per sample

Additional file 1, Supplementary Table 4: All SBS signature contributions, indels counts, and SV counts, per sample, log-transformed and scaled to z-scores

Additional file 3: Supplementary text; pdf (.pdf)

Supplementary Text: Significant models that are not described in the main text

Additional file 4: Supplementary figures; pdf (.pdf)

Supplementary Fig. 1: Shortlisted predictive models of DDR gene deficiencies not included in the main manuscript

Supplementary Fig. 2: *CDKN2A*-d in PCAWG and HMF skin cancers, *HERC2*-d in HMF skin cancers, and *ARID1A*-d in PCAWG prostate cancers

## REFERENCE LIST

1. Lindahl, T. Instability and decay of the primary structure of DNA. Nature 362, 709–715 (1993).

2. Nik-Zainal, S. et al. Mutational Processes Molding the Genomes of 21 Breast Cancers. Cell vol. 149 979–993 (2012).

3. Volkova, N. V. et al. Mutational signatures are jointly shaped by DNA damage and repair. Nat. Commun. 11, 2169 (2020).

4. Bryant, H. E. et al. Specific killing of BRCA2-deficient tumours with inhibitors of poly(ADP- ribose) polymerase. Nature vol. 447 346–346 (2007).

5. Le, D. T. et al. PD-1 Blockade in Tumors with Mismatch-Repair Deficiency. N. Engl. J. Med. 372, 2509–2520 (2015).

6. von Bueren, A. O. et al. Mismatch repair deficiency: a temozolomide resistance factor in medulloblastoma cell lines that is uncommon in primary medulloblastoma tumours. Br. J. Cancer 107, 1399–1408 (2012).

7. Paculová, H. et al. BRCA1 or CDK12 loss sensitizes cells to CHK1 inhibitors. Tumour Biol. 39, 1010428317727479 (2017).

8. Landrum, M. J. et al. ClinVar: public archive of relationships among sequence variation and human phenotype. Nucleic Acids Res. 42, D980–5 (2014).

9. Davies, H. et al. HRDetect is a predictor of BRCA1 and BRCA2 deficiency based on mutational signatures. Nat. Med. 23, 517–525 (2017).

10. Nguyen, L., W M Martens, J., Van Hoeck, A. & Cuppen, E. Pan-cancer landscape of homologous recombination deficiency. Nat. Commun. 11, 5584 (2020).

11. McVey, M. & Lee, S. E. MMEJ repair of double-strand breaks (director’s cut): deleted sequences and alternative endings. Trends Genet. 24, 529–538 (2008).

12. Nussenzweig, A. & Nussenzweig, M. C. A backup DNA repair pathway moves to the forefront. Cell 131, 223–225 (2007).

13. Umar, A. et al. Defective mismatch repair in extracts of colorectal and endometrial cancer cell lines exhibiting microsatellite instability. J. Biol. Chem. 269, 14367–14370 (1994).

14. Edelmann, W. et al. The DNA mismatch repair genes Msh3 and Msh6 cooperate in intestinal tumor suppression. Cancer Res. 60, 803–807 (2000).

15. Lanni, J. S. & Jacks, T. Characterization of the p53-dependent postmitotic checkpoint following spindle disruption. Mol. Cell. Biol. 18, 1055–1064 (1998).

16. Gorgoulis, V. G. et al. Activation of the DNA damage checkpoint and genomic instability in human precancerous lesions. Nature 434, 907–913 (2005).

17. Popova, T. et al. Ovarian Cancers Harboring Inactivating Mutations in CDK12 Display a Distinct Genomic Instability Pattern Characterized by Large Tandem Duplications. Cancer Res. 76, 1882–1891 (2016).

18. Menghi, F. et al. The Tandem Duplicator Phenotype Is a Prevalent Genome-Wide Cancer Configuration Driven by Distinct Gene Mutations. Cancer Cell 34, 197–210.e5 (2018).

19. 19. ICGC/TCGA Pan-Cancer Analysis of Whole Genomes Consortium. Pan-cancer analysis of whole genomes. Nature 578, 82–93 (2020).

20. Priestley, P. et al. Pan-cancer whole-genome analyses of metastatic solid tumours. Nature 575, 210–216 (2019).

21. Tate, J. G., et al. COSMIC: the Catalogue Of Somatic Mutations In Cancer. Nucleic Acids Res. 47, D941–D947 (2019).

22. Alexandrov, L. B. & Stratton, M. R. Mutational signatures: the patterns of somatic mutations hidden in cancer genomes. Curr. Opin. Genet. Dev. 24, 52–60 (2014).

23. Pearl, L. H., Schierz, A. C., Ward, S. E., Al-Lazikani, B. & Pearl, F. M. G. Therapeutic opportunities within the DNA damage response. Nat. Rev. Cancer 15, 166–180 (2015).

24. Knijnenburg, T. A. et al. Genomic and Molecular Landscape of DNA Damage Repair Deficiency across The Cancer Genome Atlas. Cell Rep. 23, 239–254.e6 (2018).

25. Olivieri, M. et al. A Genetic Map of the Response to DNA Damage in Human Cells. Cell 182, 481–496.e21 (2020).

26. Rentzsch, P., Witten, D., Cooper, G. M., Shendure, J. & Kircher, M. CADD: predicting the deleteriousness of variants throughout the human genome. Nucleic Acids Research vol. 47 D886–D894 (2019).

27. Degasperi, A. et al. A practical framework and online tool for mutational signature analyses show inter-tissue variation and driver dependencies. Nat Cancer 1, 249–263 (2020).

28. Davis, J. & Goadrich, M. The relationship between Precision-Recall and ROC curves. in Proceedings of the 23rd international conference on Machine learning 233–240 (Association for Computing Machinery, 2006).

29. Ceccaldi, R. et al. Homologous-recombination-deficient tumours are dependent on Polθ- mediated repair. Nature 518, 258–262 (2015).

30. Hanel, W. & Moll, U. M. Links between mutant p53 and genomic instability. J. Cell. Biochem. 113, 433–439 (2012).

31. Graham, L. S. et al. Mismatch repair deficiency in metastatic prostate cancer: Response to PD-1 blockade and standard therapies. PLoS One 15, e0233260 (2020).

32. Blazek, D. et al. The Cyclin K/Cdk12 complex maintains genomic stability via regulation of expression of DNA damage response genes. Genes Dev. 25, 2158–2172 (2011).

33. Marqués, F. et al. A new subfamily of high molecular mass CDC2-related kinases with PITAI/VRE motifs. Biochem. Biophys. Res. Commun. 279, 832–837 (2000).

34. Li, X., Chatterjee, N., Spirohn, K., Boutros, M. & Bohmann, D. Cdk12 Is A Gene-Selective RNA Polymerase II Kinase That Regulates a Subset of the Transcriptome, Including Nrf2 Target Genes. Sci. Rep. 6, 21455 (2016).

35. Li, Y. et al. Patterns of somatic structural variation in human cancer genomes. Nature 578, 112–121 (2020).

36. Wu, Y.-M. et al. Inactivation of CDK12 Delineates a Distinct Immunogenic Class of Advanced Prostate Cancer. Cell vol. 173 1770–1782.e14 (2018).

37. Rescigno, P. et al. Characterizing CDK12-Mutated Prostate Cancers. Clinical cancer research: an official journal of the American Association for Cancer Research vol. 27 566– 574 (2021).

38. Sumanasuriya, S. & De Bono, J. Treatment of Advanced Prostate Cancer—A Review of Current Therapies and Future Promise. Cold Spring Harb. Perspect. Med. 8, (2018).

39. Bajrami, I. et al. Genome-wide profiling of genetic synthetic lethality identifies CDK12 as a novel determinant of PARP1/2 inhibitor sensitivity. Cancer Res. 74, 287–297 (2014).

40. Joshi, P. M., Sutor, S. L., Huntoon, C. J. & Karnitz, L. M. Ovarian cancer-associated mutations disable catalytic activity of CDK12, a kinase that promotes homologous recombination repair and resistance to cisplatin and poly(ADP-ribose) polymerase inhibitors. J. Biol. Chem. 289, 9247–9253 (2014).

41. Mukherjee, J. et al. Mutant IDH1 Cooperates with ATRX Loss to Drive the Alternative Lengthening of Telomere Phenotype in Glioma. Cancer Res. 78, 2966–2977 (2018).

42. Alexandrov, L. B. et al. The repertoire of mutational signatures in human cancer. Nature 578, 94–101 (2020).

43. Singh, V. K., Rastogi, A., Hu, X., Wang, Y. & De, S. Mutational signature SBS8 predominantly arises due to late replication errors in cancer. Commun Biol 3, 421 (2020).

44. Alexandrov, L. B. et al. Clock-like mutational processes in human somatic cells. Nat. Genet. 47, 1402–1407 (2015).

45. Alexandrov, L. B. et al. Signatures of mutational processes in human cancer. Nature vol. 500 415–421 (2013).

46. Nik-Zainal, S. et al. The genome as a record of environmental exposure. Mutagenesis gev073 (2015) doi:10.1093/mutage/gev073.

47. Sauter, J. L. et al. SMARCA4-deficient thoracic sarcoma: a distinctive clinicopathological entity with undifferentiated rhabdoid morphology and aggressive behavior. Mod. Pathol. 30, 1422–1432 (2017).

48. Rekhtman, N. et al. SMARCA4-Deficient Thoracic Sarcomatoid Tumors Represent Primarily Smoking-Related Undifferentiated Carcinomas Rather Than Primary Thoracic Sarcomas. J. Thorac. Oncol. 15, 231–247 (2020).

49. Dagogo-Jack, I. et al. Clinicopathologic Characteristics of BRG1-Deficient NSCLC. J. Thorac. Oncol. 15, 766–776 (2020).

50. Concepcion, C. P. et al. SMARCA4 inactivation promotes lineage-specific transformation and early metastatic features in the lung. Cancer Discov. (2021) doi:10.1158/2159-8290.CD-21-0248.

51. Liu, L. et al. Mutation of the CDKN2A 5’UTR creates an aberrant initiation codon and predisposes to melanoma. Nat. Genet. 21, 128–132 (1999).

52. Amos, C. I. et al. Genome-wide association study identifies novel loci predisposing to cutaneous melanoma. Hum. Mol. Genet. 20, 5012–5023 (2011).

53. Cubillos-Rojas, M. et al. The E3 ubiquitin protein ligase HERC2 modulates the activity of tumor protein p53 by regulating its oligomerization. J. Biol. Chem. 289, 14782–14795 (2014).

54. Li, J. et al. PTEN, a putative protein tyrosine phosphatase gene mutated in human brain, breast, and prostate cancer. Science 275, 1943–1947 (1997).

55. Liaw, D. et al. Germline mutations of the PTEN gene in Cowden disease, an inherited breast and thyroid cancer syndrome. Nat. Genet. 16, 64–67 (1997).

56. Fda. FDA approves niraparib for HRD-positive advanced ovarian cancer. (2019).

57. Boland, C. R. & Goel, A. Microsatellite instability in colorectal cancer. Gastroenterology 138, 2073–2087.e3 (2010).

58. Umar, A. et al. Revised Bethesda Guidelines for hereditary nonpolyposis colorectal cancer (Lynch syndrome) and microsatellite instability. J. Natl. Cancer Inst. 96, 261–268 (2004).

59. Abida, W. et al. Analysis of the Prevalence of Microsatellite Instability in Prostate Cancer and Response to Immune Checkpoint Blockade. JAMA Oncol 5, 471–478 (2019).

60. Antonarakis, E. S. et al. Clinical Features and Therapeutic Outcomes in Men with Advanced Prostate Cancer and DNA Mismatch Repair Gene Mutations. Eur. Urol. 75, 378– 382 (2019).

61. Xue, Y. et al. SMARCA4 loss is synthetic lethal with CDK4/6 inhibition in non-small cell lung cancer. Nat. Commun. 10, 557 (2019).

62. Malcikova, J. et al. Monoallelic and biallelic inactivation of TP53 gene in chronic lymphocytic leukemia: selection, impact on survival, and response to DNA damage. Blood 114, 5307–5314 (2009).

63. Evers, B. & Jonkers, J. Mouse models of BRCA1 and BRCA2 deficiency: past lessons, current understanding and future prospects. Oncogene 25, 5885–5897 (2006).

64. Reitmair, A. H. et al. MSH2 deficient mice are viable and susceptible to lymphoid tumours. Nat. Genet. 11, 64–70 (1995).

65. Prolla, T. A. et al. Tumour susceptibility and spontaneous mutation in mice deficient in Mlh1, Pms1 and Pms2 DMA mismatch repair. Nat. Genet. 18, 276–279 (1998).

66. Donehower, L. A. The p53-deficient mouse: a model for basic and applied cancer studies. Semin. Cancer Biol. 7, 269–278 (1996).

67. Karczewski, K. J. et al. The mutational constraint spectrum quantified from variation in 141,456 humans. Nature 581, 434–443 (2020).

68. Itan, Y. et al. The mutation significance cutoff: gene-level thresholds for variant predictions. Nat. Methods 13, 109–110 (2016).

69. Nik-Zainal, S. et al. Landscape of somatic mutations in 560 breast cancer whole-genome sequences. Nature 534, 47–54 (2016).

70. Friedman, J., Hastie, T. & Tibshirani, R. Regularization paths for generalized linear models via coordinate descent. J. Stat. Softw. (2010).

