## Additional file 3; Supplementary text for "Pan-cancer association of DNA repair deficiencies with whole-genome mutational patterns"

### Significant models that are not described in the main text

**Monoallelic predictive model of *ARID1A*-d in HMF prostate cancer patients**

Loss of *ARID1A* impairs the pausing of RNA polymerase II doing transcription, leading to dys-regulated expression of genes(63). We predicted *ARID1A* LOF (PR-AUC-E=0.208; AUROC=0.72) by depletion of SBS signature 8 mutations(**Additional file 4, Supp. Fig. 1a; Additional file 1, Supp. Table 7**). SBS signature 8 mutations have been associated with inactivity of *BRCA1/2(8)* and we found that the number of SBS signature 8 mutations, compared to prostate cancers which are wild type in both *ARID1A* and *BRCA1l2*, was significantly depleted (Wilcox. Rank-sum test $p<0.0013$). As expected, the difference became much more pronounced compared to *BRCA1/2* mutated prostate cancers ($p<7.7x10^{-5}$) (**Additional file 4, Supp. Figure 2d**). This suggests differences in cancer evolution between prostate cancers with and without *ARID1A*-d as well as those that are homologous recombination deficient.

***BAP1* associated with increased mutational SBS signature 30 (defective BER) and decreased SBS signature 7 in HMF skin cancers**

*BAP1* germline variants are commonly associated with predisposition for the development of multiple cancer types including melanoma(64). We could predict the deficiency of *BAP1* (PR-AUC-E=0.26; AUROC=0.80) primarily by a decreased number of SBS signature 7 mutations and an increase number of SBS signature 30 mutations (**Additional file 4, Supp. Fig. 1b; Additional file 1, Supp. Table 7**). The UV-induced SBS signature 7 (**Additional file 1, Supp. Table 5**) is generally dominant in skin cancers(65,66) and SBS signature 30 which is associated with inefficient base excision repair (67) (**Additional file 1, Supp. Table 5**). SBS signature 30 has a high compositional similarity to SBS signature 7a (cosine similarity=0.72). The co-occurrence and opposite predictive effects of the two signatures may thus be an affect of technical difficulties with their combined inference.

#### ***MEN1*** **deficiency is associated with depletion of SBS signature 9 (POLH hypermutation activity) as well as SBS signature 16 (unknown etiology)**

Ten out of a hundred neuro-endocrine cancers (NET; HMF) have biallelic loss of the *MEN1* gene, with systematically fewer mutations attributable to signature 9, which is related to hyperactivity of POLH (PR-AUC-E=0.22; AUROC=0.82) (**Additional file 4, Supp. Fig. 1c; Additional file 1, Supp. Table 7**). *MEN1* is a regulator of gene transcription and germline deficiencies are causatively associated with developing Multiple Endocrine Neoplasia Type 1 (MEN1), which is a rare, hereditary tumor condition(68). However, in this setting, we observed only somatic events, with all ten patients having a LOH combined with a pathogenic, somatic variants: six variants were insertions or deletions in the coding region, three were stop-gain SBS, and a single was a splice-site variant (**Additional file 1, Supp. Table 8**). This agrees with studies of somatic *MEN1* mutation that likewise found somatic hits in *MEN1* exclusively together with LOH events, and associated the somatic LOF of *MEN1* with a different disease phenotype than that of inherited MEN1(69).

#### ***RB1*** **LOF associated with SBS signature 7 in cancers of the urinary tract**

Across the 21 biallelic *RB1*-d HMF urinary tract cancers (n=108), we observed an increased number of mutations attributable to UV-related SBS signature 7 (PR-AUC=0.47; AUROC=0.84) (**Additional file 4, Supp. Fig. 1d; Additional file 1, Supp. Table 7**). SBS signature 7 is known to develop from the exposure to UV light (**Additional file 1, Supp. Table 5**)(66). However the signature has been reported by COSMIC for various cancer types with no sun exposure, including cancers of the breast, ovary, pancreas, oral cavity, lung, and uterus as well as sarcomas(17). Although the predictive performance is considerable and significant, the accumulation of mutations in SBS signature 7 does not necessarily reveal the true etiology of the underlying mutagenesis.
