## Additional file 4; Supplementary figures for "Pan-cancer association of DNA repair deficiencies with whole-genome mutational patterns"

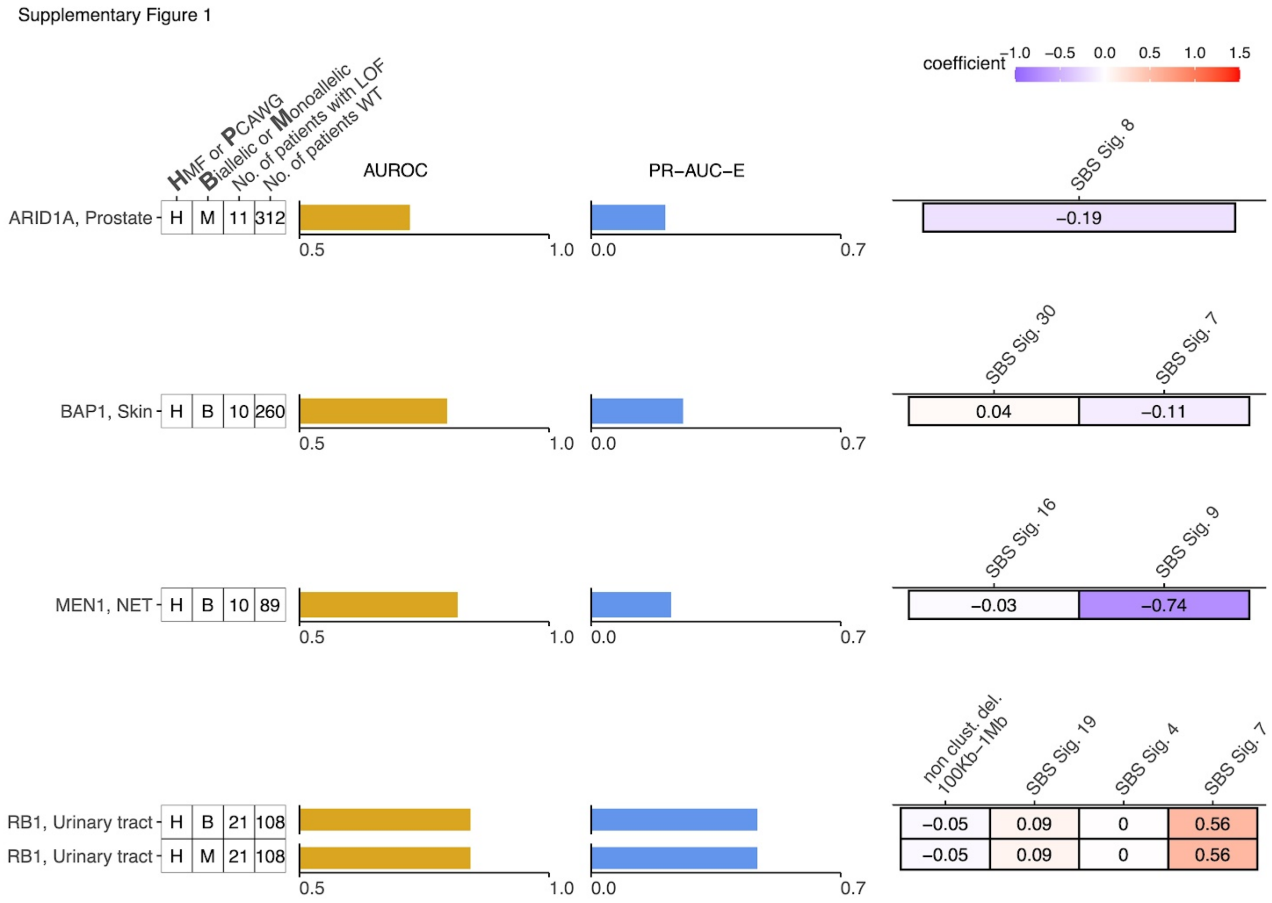


**Supp. Fig. 1: Shortlisted predictive models of DDR gene deficiencies not included in the main manuscript**

**a** Predictive model of *ARID1A* with Precision-Recall area-under-the-curve enrichment from the baseline (baseline - rate of LOF in the data set; PR-AUC-E) and area-under-the-roc curve (AUROC), as well as feature coefficients. Same info for predictive models of **b** *BAP1,* **c** *MEN1*, and **d** *RB1*

Supplementary Figure 2

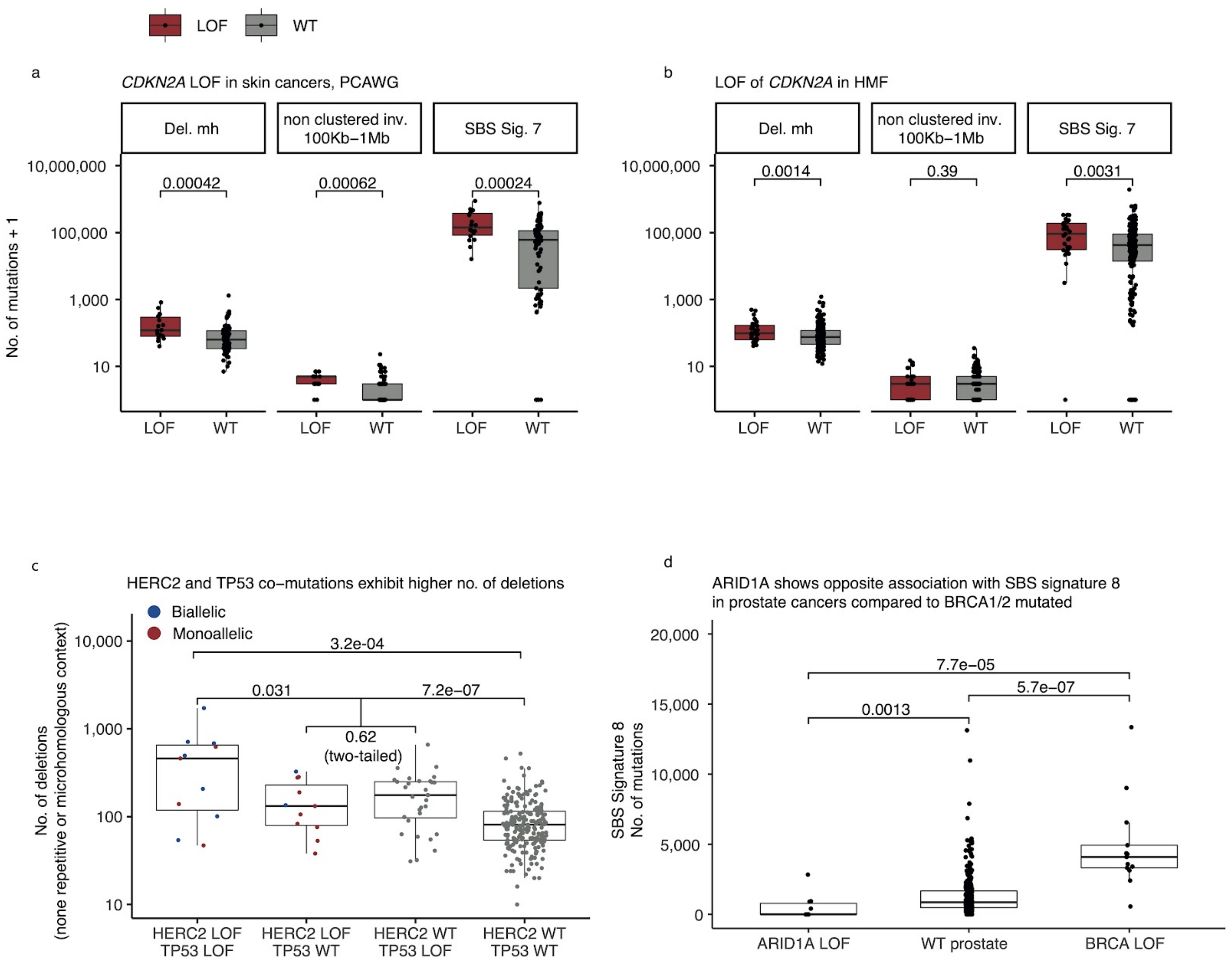


**Supp. Fig. 2: *CDKN2A*-d in PCAWG and HMF skin cancers, *HERC2*-d in HMF skin cancers, and *ARID1A*-d in PCAWG prostate cancers**

**a** *CDKN2A* LOF cancers (red) and wild-type (grey) compared by the number of mutations (y-axis; logarithmic) across patients from the PCAWG and **b** the HMF data set (wilcoxon rank-sum test, one-tailed). **c** Number of deletions in HMF skin cancers, *HERC2*-d (biallelic LOF red; monoallelic blue) and monoallelic *TP53*-d compared to patients with LOF of either gene (Wilcox.

Test one-tailed), and to patients with LOF in neither gene (Wilcox. Test one-tailed). Patients with mutations in either *TP53* or *HERC2* are compared (Wilcox. Test two-tailed). Comparison of patients with LOF of both genes and patients with exclusively *TP53* LOF (not included; Wilcox. Test one-tailed p=0.049)**. d** Number of SBS sig. 8 mutations in *ARID1A* mutated prostate cancers, *ARID1A* wild-type cancers, and *BRCA1/2* mutated prostate cancers (Wilcox. Test, one-tailed).
